# From Criticism to Collaboration: Empowering Built-Environment Professionals for Biodiversity-inclusive Cities in the Global North and South

**DOI:** 10.1101/2025.01.15.633110

**Authors:** Jing Lu, Li Li, Ana Hernando, Jose Antonio Manzanera

## Abstract

Cities play a crucial role in addressing the biodiversity crisis. Despite advancements in urban ecology, effectively integrating biodiversity into urban planning remains limited. Built-environment professionals (BEPs), who are responsible for local actions, often face criticism for their unwillingness to prioritize biodiversity and are seldom studied from their perspectives, especially those from the Global South. Therefore, understanding the viewpoints of diverse BEPs from both the Global South and North on biodiversity-inclusive urban planning and design (BIPD) is crucial. This study employs Q-Methodology to analyze BEPs’ perspectives in Ethiopia, China, Spain, and the United States, revealing four viewpoints: Pluralist, Enthusiast, Pragmatist, and Contextualist. Challenges in realizing BIPD include widespread unawareness among BEPs of existing biodiversity data and methodologies, structural-physical barriers such as land use conflicts and competing resources, and misaligned values with stakeholders. Our research challenges the notion that BEPs resist incorporating biodiversity into their practices by broadly exploring their viewpoints. BEPs exhibit readiness to transcend legislative structural barriers and leverage professional norms to advocate for urban biodiversity, particularly when equipped with evidence-based tools. Pluralists and Enthusiasts demonstrate heightened interest in multidisciplinary collaboration and prescriptive, visual tools to communicate biodiversity’s value to stakeholders. Critically, BEPs’ capacity to act hinges on the availability of contextually sensitive methods, necessitating empowerment through transdisciplinary collaborations to foster transformative changes in cities, especially in the Global South, where complex socioeconomic-ecological trade-offs demand co-produced solutions.

## Introduction

Cities increasingly play a vital role in global biodiversity conservation and enhancement^1,2,3^, There is a growing recognition of the significance of urban biodiversity, as it contributes to countering the global extinction crisis^2,4^, promoting multispecies justice^5,6^, and providing synergetic benefits for human well-being^7,8,9^ as well as climate change mitigation and adaptation^10,11^. Despite these recognitions, biodiversity in cities continues to decline, impacting even common species such as the house sparrow^12^. To turn these recognitions into real-world actions, innovations in urban planning and design practices that enhance native biodiversity and support inclusive and sustainable urbanization are essential. This concept has been acknowledged by several global targets, initiatives, and commitments^13,14^. It is encapsulated as biodiversity-inclusive urban planning and design (BIPD), a term coined and promoted in the Kunming-Montreal Global Biodiversity Framework (KMGBF) adopted by the Convention on Biological Diversity^15^.

Progress towards such global initiatives is rooted in local actions^14^. However, urban professionals in the built environment responsible for engaging citizens in local decision-making may not yet be prepared for BIPD. Built-environment professionals (BEPs), such as urban planners, civil engineers, and landscape architects, have historically focused on improving human well-being rather than considering the needs of other organisms and often lack a background in ecological education^12^. Such a situation is more challenging in the Global South, where more generalists lack the capacity for specialization^16^, and more attention should be directed toward citizens’ basic needs^17^. Although there is a long tradition of integrating ecological principles into the planning process, guided by landscape urbanism^18,19^, and movements centered on concepts and practices such as urban green infrastructure planning^20^ and biophilic urbanism^21^, BEPs’ practices have faced criticism from ecologists. Concerns include a focus on the expansion and the connectivity of green spaces and neglecting the potential of the built environment as habitat^22^, treating biodiversity largely as incidental and passive stakeholders rather than engaging actively and explicitly^23^, a relative lack of attention toward animals^24^, and oversimplifying the complexity of ecological relationships and their interactions^12^.In this regard, BEPs are questioned about their willingness and ability to navigate the complexities of ecological factors intertwined with the socioeconomic and political intricacies of planning processes^16,24,25^.

While BEPs are being questioned, the shift in urban ecology from a biologically based discipline to an increasingly interdisciplinary field^27,28,29,30^ offers promising opportunities to advance knowledge necessary to explicitly involve urban biodiversity as nonhuman stakeholders in planning and decision-making^23^. Recent advancements in urban ecology have produced sophisticated knowledge and tools that have the potential to inform urban planning and design. For instance, Plummer et al. developed species-habitat models to predict bird populations for urban development planning scenarios, aiming to support the on-site compensation policy of “no biodiversity net loss” in the UK^32^. By employing citizen-science data on animal species and Maximum entropy (MaxEnt) species distribution modeling (SDM), researchers in China and South Korea have mapped habitat cores and ecological corridors for urban biodiversity conservation^33,34^. Furthermore, recent efforts in urban ecology have demonstrated the potential for actively integrating biodiversity science into urban practices. Building on the target species selection framework^22^ and species’ life-cycle study^35^, Weisser, Hauck, and their teams^12^ employed the Animal-Aided Design approach to explicitly consider animals as stakeholders in their project design practices in Germany and Switzerland. Guided by the framework of Biodiversity Sensitive Urban Design^36^. Australia’s largest urban renewal project, Fishermans Bend in Melbourne, stands out as one of the earliest urban developments of this scale to explicitly incorporate biodiversity targets^37^.

Nevertheless, these advancements in urban ecology have not been widely reflected in practical urban planning and design. To date, 66% of research on urban planning and design focusing on urban biodiversity is still guided by the idea that “urban form leads to changes in species diversity within urban areas,” treating nonhuman species as passive stakeholders^23^. Although limited in number, studies have attempted to investigate the reasons and challenges behind this scenario. Some studies attribute this phenomenon to BEPs’ value bias and lack of willingness to prioritize biodiversity compared to other competing interests^38,39^. Additionally, knowledge gaps and disciplinary silos are discussed, emphasizing BEPs’ lack of empirical ecological knowledge^16^ necessary for prescribing “hands-on” transdisciplinary real-world solutions^16,26,40^. Moreover, BEPs may experience path dependence^41^ and are considered to rely on their socio-technical regimes, leading to resistance to change or reluctance to seek external collaborations^42,43^. While these studies provide an initial understanding of the challenges, they tend to critique BEPs without thoroughly examining their perspectives on BIPD. They engage only a specific interest group, including some BEPs attracted to the Summit related to urban wildlife^39^, excluding most BEPs that shape the urban landscape, which usually focuses on various social, economic, and environmental objectives. Traditionally, few BEPs engage with biodiversity topics, as biodiversity is not an explicit goal in their practices but rather a constraint that must be managed when necessary^12^. Furthermore, studies involving BEPs mostly concentrate on related topics, such as Nature-based Solutions (NbS)^42^, which do not specifically address BIPD; instead, the value of urban biodiversity is often overlooked in the design and assessment of many NbS programs^1^. Even when considering studies on related topics, the voices of BEPs from the Global South are severely underrepresented despite their pressing urban challenges amidst limited resources^16^. Without specifically understanding the perspectives of a diverse group of BEPs from various professional backgrounds and global regions regarding BIPD, it is difficult to transform global biodiversity commitments from being merely advocated by ecologists^16^ to fostering real-world actions. Additionally, these discussions often stop at identifying barriers^39,42^ rather than exploring opportunities involving BEPs.

To fill this gap, this study investigates the perspectives of a broad profile of BEPs across four countries in both the Global South and North regarding BIPD, drawing on theories of Lock-in Effects^41^ and the Theory of Planned Behavior (TPB)^44^. The TPB is one of the most influential models for predicting and explaining individuals’ intention to engage in specific behaviors^45^. It deconstructs a person’s behavioral intentions into three constructs: attitudes, social and subjective norms, and perceived behavioral control^44^. Recently, some studies have extended the TPB model by incorporating additional constructs, such as moral norms to understand green purchasing behavior^45^ and personal norms alongside environmental concerns for public transit behavioral intentions^46^. By extensively comparing various extensions of TPB research, these studies argue that moral and environmental concerns should be included as an extension of TPB, which are significant predictors of intentions on environmentally responsible behaviors^44,46^. In addition, Wilson^41^ discussed how path dependency and “lock-in” effects pose severe hindrances to social-ecological resilience and transitional changes, as they can trap stakeholders in pathways from which they find it difficult to escape. He categorized lock-in effects at the community level into three types: 1) structural, which are beyond the control of individual communities; 2) economic, related to communities’ economic capital; and 3) social-psychological, associated with community-level endogenous social and psychological factors^41^. Given the potential challenges of implementing BIPD, it is appropriate to reference the extension of TPB and Lock-in effects as an initial framework to help understand the BEPs’ intentions in involving biodiversity as nonhuman stakeholders in their practice, specifically taking citizen-science-based species distribution modeling (SDM), which is widely studied in urban ecology^31,34^, as an example. We aim to address three knowledge gaps: 1) uncovering the diverse perspectives of various BEPs’ regarding BIPD, 2) providing in-depth, specific, and nuanced evidence of the perspectives of various BEPs across the Global North and South to extend to those understudied and hard to reach, and 3) understanding the challenges and opportunities of implementing BIPD through the lens of BEPs in relation to the frameworks of the TPB and Lock-in effects.

## Results

We explored the perspectives of BEPs using the standard protocols of Q-Methodology, which involved two stages of interviews: the Concourse and Q-sets. This was conducted with a carefully selected group of case study countries from both the South and North, along with purposively sampled BEPs to ensure diversity (Table 1). During the Concourse and Q sorts interviews, we discovered that only four out of the 36 participants had heard or used SDMs before, representing just 11.1% of the BEPs surveyed. We identified four factors (F) that account for 19 of the 23 participants’ Q sorts and revealed four distinct viewpoints on BIPD (Table 2). The remaining four Q sorts were classified as confounded Q sorts (Table 2) due to significant loading onto multiple factors. We named the four factors *Pluralist, Enthusiast, Pragmatist*, and *Contextualist*. Notably, all perspectives support BIPD and are independent of participants’ countries or professional backgrounds (Table 1 and 2). Professionals from Ethiopia appear in all factors, professionals from the United States in three factors, and professionals from China and Spain in two factors. Similarly, there is no clear pattern among participants’ professional backgrounds, as they are also distributed across factors.

**Table 1:** Summary of research participants

| Study country |  |  | Professional background |  |  |  |  |  |  |  |  | Total participants per country |  |
| --- | --- | --- | --- | --- | --- | --- | --- | --- | --- | --- | --- | --- | --- |
| Name | National Urbanization (%) | National GDP per capita, (PPP) (current international \$) | Discipline | | | | | Focus | | | | Concourse | Q-sort |
|  |  |  | UP | UD | LA | CE | Env | Mul | Ecol | Environ | Socio-econ |  |  |
| Ethiopia | Low | Low | 4 | 1 | 1 | 4 | 0 | 0 | 0 | 5 | 5 | 4 | 6 |
| China | Medium | Lower middle | 3 | 2 | 1 | 0 | 2 | 1 | 2 | 2 | 5 | 3 | 6 |
| Spain | High | Upper middle | 1 | 1 | 2 | 3 | 1 | 0 | 2 | 3 | 3 | 3 | 5 |
| USA | High | High | 3 | 0 | 2 | 1 | 1 | 2 | 3 | 3 | 3 | 3 | 6 |
| Total participants per background |  |  | 11 | 4 | 6 | 8 | 4 | 3 | 7 | 13 | 16 | 13 | 23 |

Source: National urbanization (urban population as a percentage of total population) (%): Low < 35%, Medium 35%-75%, High > 75% <sup>47</sup>; National GDP per capita, purchasing power parity (PPP) (current international \$): Low < 15, Lower middle 15-45, Upper middle 45-65, High > 65<sup>48</sup>.
UP = urban planning; UD = urban design; LA = landscape architecture; CE = civil engineer; Env = environmental science or engineering; Mul = specialized in two or more disciplines; Ecol = ecological focus; Environ = environmental focus; Socio-econ = socioeconomic focus.

**Table 2:**
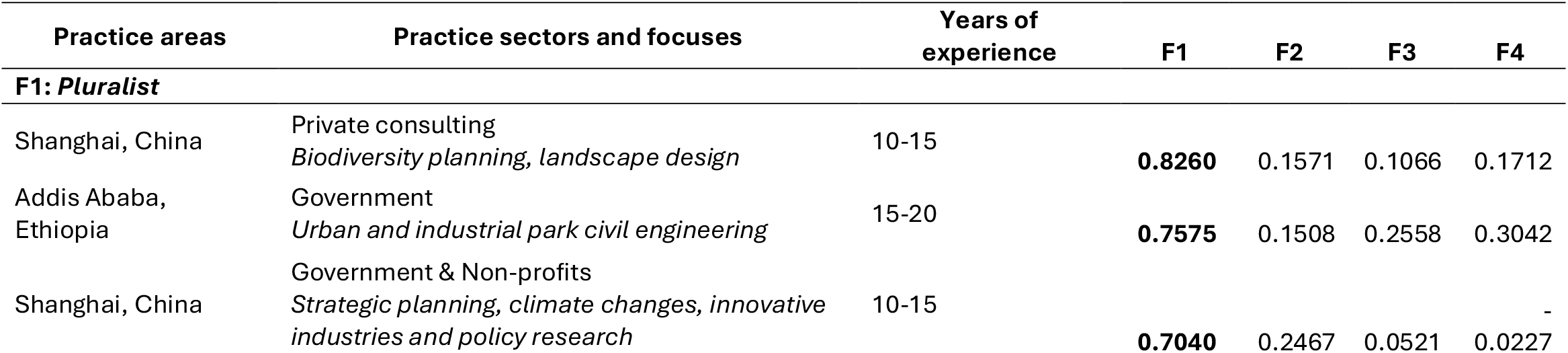
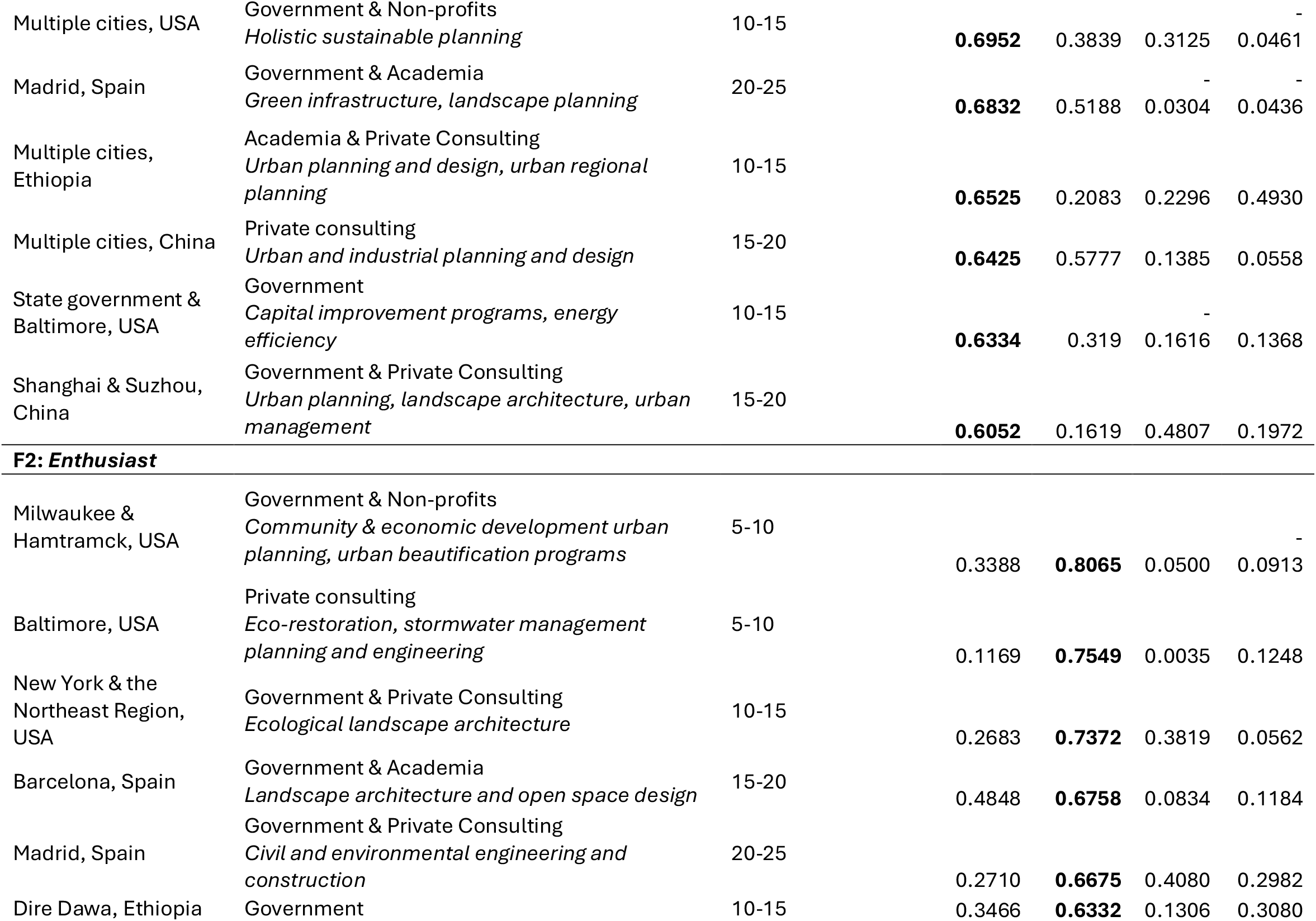
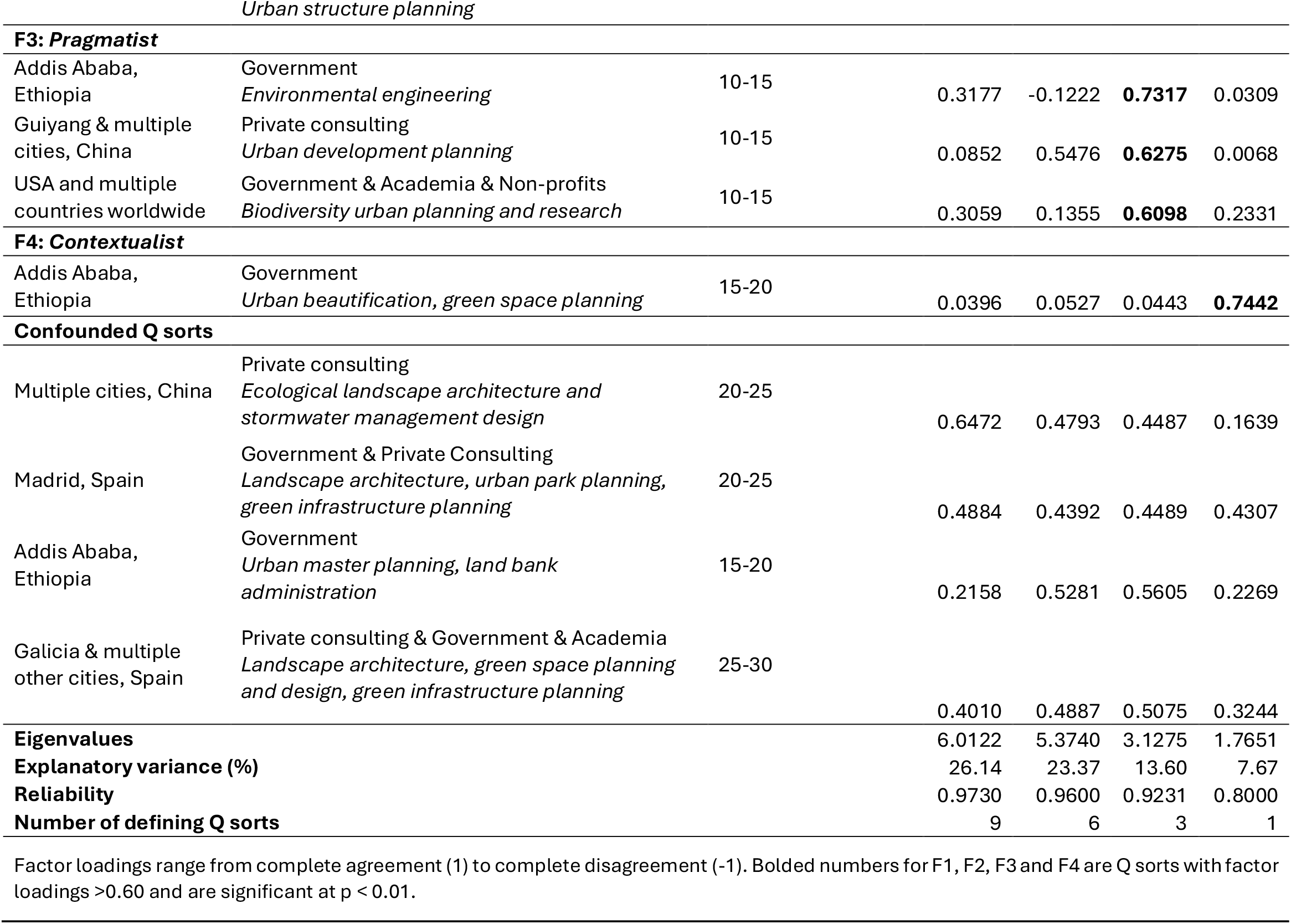
Factor loadings for each participant’s Q sort

### 3.1 Factor 1 – Pluralist

The Pluralist views BIPD holistically, recognizing the importance of biodiversity interwoven with many other public interests and considering how to balance them together. This perspective is shared by the largest number of participants across countries and backgrounds. They regard biodiversity as an interwoven fabric in a holistic system and cannot be discussed separately from other socioeconomic factors. In this context, they feel neutral about prioritizing socioeconomic needs in urban practice (S9, 0) (Table 3). As one participant from China said: “*It frames in a way that excludes biodiversity from the societal socioeconomic needs. But they are interconnected, and the ecological benefits of the urban socioeconomic development*.” They are also neutral about the statement “BIPD has to be biodiversity centric, as the earth is increasingly losing biodiversity to a tipping point (S1, 0**)”, while all other viewpoints support this statement. They do not relate to the urgency and instead adopt a contextually-sensitive approach. One professional from China noted: “*It depends on the nature of the projects. Some projects may prioritize biodiversity as it fits with the contextual situation and project goal…Then planners may need to balance socioeconomic requirements with biodiversity*.” They slightly disagree (S20, -1**) with the statement that it is luxurious to consider the needs of non-human species. As one participant from the United States stated, “*On a practical level, it is really challenging to balance providing for other species when there are vulnerable humans whose needs should be met*.” One option to balance the competing interests is through time: “*It almost feels like a short-term versus a long-term issue. We must prioritize meeting the needs of the people who are in vulnerable states now*.”

**Table 3:** Q Statements and rankings for each factor

| **Q Statements** | **F 1** | **F2** | **F 3** | **F 4** |
| --- | --- | --- | --- | --- |
| 1. BIPD has to be biodiversity centric, as the earth is increasingly losing biodiversity to a tipping point | 0** | 2* | 4 | 4 |
| 2. Humans should conserve biodiversity because it is essential to our mental health | 2 | 4 | 3 | 2** |
| 3. The most important reason for biodiversity conservation is human survival | 1 | 1 | 3** | -1 |
| 4. BIPD is a valuable layer in considering development patterns and how to use space | 3 | 2 | 1 | 1 |
| 5. BIPD gets down to a granular enough detail that are useful for the urban practice | 2* | 0 | 1 | 0 |
| 6. It is very likely to mainstream BIPD into the urban practice in the near future | 3 | 1** | 2 | 2 |
| 7. In the urban settings, BIPD is a suitable approach to reduce human disturbance to the natural habitats | 2 | 2 | 1** | 3 |
| 8. BIPD helps represent non-human species' voices in the urban practice | 4* | 1 | -1 | 0 |
| 9. Built environment professionals have to prioritize socioeconomic needs before considering biodiversity in the urban practice | 0 | -2 | 2* | -1 |
| 10. As built environment professionals, we should not leave out biodiversity information or data if they are available** | 1 | 3 | 2 | 0 |
| 11. It is not necessary for built environment professionals to involve BIPD in the urban practice because there are no such mandatory regulations | -4 | -4 | 0 | 0 |
| 12. The primary challenge for BIPD is that built environmental professionals do not have quality non-human species data in cities to inform practice* | 0 | -1 | 0 | 1 |
| 13. The major barrier to involve BIPD into urban practice is that built environment professionals do not know how to do it | -1 | 0 | 1 | -1 |
| 14. Despite understanding differences, built environment professionals can still work effectively with ecologists and biologists in the BIPD projects | 1 | 1 | 1 | 3** |
| 15. It is not practical to involve BIPD in the urban practice because biodiversity knowledge is too specific and narrowly focused | 3** | -1 | 0 | 0 |
| 16. Planning and design are not science, so biodiversity scientific data cannot be translated into design solutions or policy decisions | -3 | -2 | 0 | -1 |
| 17. As built environment professionals, we cannot make the biodiversity-inclusive shift in the urban practice because we all work in institutional silos | -2 | 0 | -2 | 1 |
| 18. It is only possible to make the biodiversity-inclusive shift in the urban practice after it is integrated into the existing regulatory framework** | 0 | -1 | 0 | 2 |
| 19. To promote BIPD, built environment professionals' major struggle is how to convey the significance and urgency of biodiversity to the stakeholders | 1 | 3 | 3** | 2 |
| 20. When humans can't meet their own needs, it's too luxurious to consider the needs of non-human species | 1** | -3 | -4 | 3** |
| 21. It is impossible to involve BIPD in cities where land is profitable and generates revenue | -1 | -3 | -3 | -2 |
| 22. The added-up of additional time, capacity and coordination efforts make it difficult to implement BIPD in the urban practice | 0 | 0 | -1 | 3** |
| 23. As built environment professionals, BIPD is not something we typically do, and we don't find the motivation to change | -2 | 0 | -2 | -2 |
| 24. In the urban settings, BIPD is only feasible in the urban expansion areas or new cities, but not in the already developed ones** | -2 | -2 | -2 | -4 |
| 25. Built environment professionals will implement BIPD projects only if there are successful examples elsewhere | -1 | 1 | -1 | 1 |

F1 =Pluralist, F2 =Enthusiast, F3 =Pragmatist and F4 =Contextualist. Statements are listed in order of the numbers on the statement cards and scores illustrate the level of agreement with each statement from -4 'strongly disagree' to +4 'strongly agree'. Distinguishing statements are noted for each factor with a \* if significant at $p < 0.05$ or \*\* if significant at $p < 0.01$ . Consensus statements are noted for each statement with a \* if non-significant at $p < 0.05$ or \*\* if non-significant at $p < 0.01$ .

Nevertheless, Pluralists value the rights of non-human species to exist and their voices to be represented, which they perceive as an objective potentially achievable through BIPD with sufficient detail. They strongly agree with the statement that “BIPD helps represent non-human species’ voices in the urban practice (S8, +4*).” A participant from the United States stated that “*we as humans have had a major impact on the Earth, and we are not the only species to live here. And we need to take other species into consideration in our plans*.” Pluralists also favor BIPD as it delves into sufficient detail (S5, +2*), leading to the following assertion from an Ethiopian professional: “*Such detailed consideration will be coming into force, and people will try to understand the impacts and consider this*.”

While expressing interest in BIPD, Pluralists also discuss their concerns about the feasibility of applying SDM examples developed by ecologists in planning practice. One professional from the United States expressed concern regarding the applicability of SDM examples, stating, “*I’m afraid that I don’t have a deep understanding of biodiversity…I love the modeling to help us make informed decisions, but I guess I would want more, like a data package to explain how to interpret them*.” Meanwhile, a participant from China, who has previously employed SDMs in her practice, further elaborated on the incompatibility of data and methods currently supported by ecologists for direct application in planning. She remarked: “*Ecology and biology are very specialized disciplines, and some experts only study one specie, but planning needs to be holistic and comprehensive. It won’t work to include only one specie in planning*.” Despite the challenges, she considered that the best way to realize BIPD is through multidisciplinary collaboration, emphasizing that some BEPs should be equipped with ecological knowledge to the extent of “*being able to effectively communicate with ecologists*. She explained, “*BEPs with a specialization in ecology are still generalists and do not need to delve deeply into scientific details, which should be the job of ecologists and biologists. But these BEPs should know how to translate and integrate these data into prescriptive planning actions*.”

This perspective also exhibits good control of their ability to go beyond mandatory regulations (S11, -4) because it works against their professional norms. One participant from the United States remarked: “*We’re not only planning because we’re required to. We’re planning to guide future development, and we have the flexibility to consider any information we need*.” This aligns with their heightened positive attitude (Table 3) regarding mainstreaming BIPD into the urban practice (S6, +3), considering BIPD an emerging hot topic and innovative direction. A professional from China described their view: “*Globally, it has become a hot theme in planning practice recently. Biodiversity is the innovative way to go for the future planning and design practice*.”

### 3.2 Factor 2 –Enthusiast

The *Enthusiast* favors the idea of BIPD and advocates prioritizing biodiversity but struggles to convey the significance to others, feeling pessimistic about mainstreaming BIPD. Professionals with this viewpoint embrace the urgency of biodiversity conservation and support incorporating biodiversity in urban practice as much as possible. A participant from the United States mentioned that “*if we are talking about incorporating biodiversity into planning and design, we should focus on what can be done to increase biodiversity. I agree with both parts of the sentence. It should be biodiversity-centric, and the Earth is increasingly losing (biodiversity) to a tipping point* (S1, +2*).” Further, they are most likely to agree not to leave out biodiversity information if available (S10, +3) (Table 3). One professional from Spain emphasized the significance of data availability of BIPD by saying: “*We should try to work with as much information and data as it is available, but especially in relation to this topic*.” Similarly, they strongly disagree with the notion of “not involving BIPD in the urban practice because there are no such mandatory regulations” (S11, -4), as articulated by a professional from Spain: “*I’m not planning just to meet mandatory regulations but to improve an existing situation*.”

Professionals sharing this perspective tend to experience a heightened sense of struggle over the prioritization of biodiversity against other considerations, with the concern that their stakeholders may not care as much. One participant from the United States remarked, “*Now the struggle mainly is to convey the urgency to others. The urgency element of environment versus other factors*. (S19, +3)” The *Enthusiast* perspective is also the only one that expresses pessimism about mainstreaming BIPD into urban practice (S6, -1*) (Table 3). A professional from the United States illustrated this with an example: “*There’s a pretty high-profile example in our city right now. What is being proposed as the revitalization of the harbor is, (in) a lot of ways, fairly similar to what’s there already. I guess that as someone who’s environmentally minded, I wish that there had been more thought into developing it as an environmental, ecological space first and foremost*.”

### 3.3 Factor 3 – Pragmatist

The *Pragmatist* emphasizes flexibility and adaptability based on real-world conditions. They view biodiversity conservation as an ongoing issue and trend, but not necessarily as an urgent one. Regarding the statement “it is very likely to mainstream BIPD into the urban practice in the near future (S6, +2)”, an Ethiopian participant expressed optimism, stating, “*because this kind of idea and issue is a (a) currently ongoing issue*.” Similarly, when talked about their strongly agreed statement that “BIPD has to be biodiversity centric, as the earth is increasingly losing biodiversity to a tipping point (S1, +4)”, one professional from China discussed that “*I think this is a core trend in our profession and practice (nowadays), and we should work toward it*.” Their interpretation does not convey a sense of urgency; rather, they view the issue through the lens of political trends or professional fashion, implying a tendency to “fit in” or “grasp opportunities.”

In contrast to other perspectives (Table 3), *Pragmatists* disagree with the assertion that “to promote BIPD, built environment professionals’ major struggle is how to convey the significance and urgency of biodiversity to their stakeholders (S19, -3**). One participant from Ethiopia indicated that “*stakeholders may be restricted to (certain) land uses due to biodiversity concerns, which would further limit a community and landowner to profit from their property*.” They appear to align with their stakeholders’ perspectives, prioritizing the latter by weighing biodiversity considerations against immediate socioeconomic concerns to arrive at more practical and tangible solutions.

*Pragmatists* justify their trade-off decisions to prioritize socioeconomic considerations by viewing biodiversity as inseparable from human well-being. Among all perspectives, this group is the most aligned with the claim that biodiversity conservation is for human survival (S3, +3**) (Table 3). A professional from the United States asserted, “*Biodiversity survival and human survival are kind of one and the same. We depend on it and when we degrade the environment, we’re degrading ourselves and our ability to eat nutritious food*.” The data from statement ranking (S9, 2*) further demonstrates their recognition of the instrumental value of biodiversity, as they uniquely align with the idea that “built environment professionals have to prioritize socioeconomic needs before considering biodiversity in urban practice” (Table 3). Their interpretation suggests that by prioritizing socioeconomic concerns, they can simultaneously address biodiversity needs, as both are interrelated. One participant from the United States emphasized, “*In the cities, we need to be able to say why biodiversity is important from an urban kind of human perspective, and if we can make those linkages between those needs, then they’ll also be considering biodiversity*.”

### 3.4 Factor 4 – Contextualist

The *Contextualist* feels a strong urgency for biodiversity conservation in the face of urbanization but is constrained by inadequate contextual capacity to achieve its goals. They underscore this urgency in response to the statement, “BIPD has to be biodiversity-centric, as the earth is increasingly losing biodiversity to a tipping point” (S1, +4). The Ethiopian professional remarked, “*Because (we are) now currently increasingly losing biodiversity at (an) alarming rate. This is a global issue now, especially in developing nations with its (rapid) urbanization and deforestation*.” They highlight the significance of biodiversity conservation in the natural habitats of developing nations to mitigate human disturbances (S7, +3): “*In the urban setting, to protect the natural habitat (away from human disturbances), biodiversity is the key issue for reducing human disturbance to the natural habitat*.*”* They primarily view humans as a threat to biodiversity within the urban context.

Despite their strong commitment to biodiversity conservation, *Contextualists* are acutely aware of their stakeholders’ struggles with primary socioeconomic needs and are attentive to the structural challenges that prevent them from realizing what they believe is right. This group uniquely agrees with the statement that “when humans can’t meet their own needs, it’s too luxurious to consider the needs of non-human species” (S20, +3**). They frame their reasoning primarily through the lens of their stakeholders, explaining: “*When human(s) cannot meet their interest or their own needs, they don’t bother about others, (especially) for developing countries that are still striving for food security, urbanization, and migration (issues)*.*”* This implies a sense of vulnerability from this professional that their stakeholders are economically locked in, while he finds it difficult to propose win-win solutions.

Meanwhile, this group perceives themselves as having a more central role in urban practice and decision-making across disciplines, which may further narrow their capacity to find effective solutions. In contrast to all other viewpoints (Table 3), they disagree with the statement that “despite understanding differences, built environment professionals can still work effectively with ecologists and biologists in the BIPD projects (14, -3**)”. They argue that “*biologists only can work (on the BIPD) unless urban planners incorporate*” despite recognizing that planning is inherently a multi-sectoral effort.

### 3.5 Consensus statements

Four statements emerged as statistically significant consensus among BEPs, with statement 12 not significant at p < 0.01, while statements 10, 18, and 24 were identified at p < 0.05 (Table 3). Among these, all BEPs across viewpoints agree only with statement 10 and disagree with statement 24. They collectively support the inclusion of biodiversity information in their practice, as “*it aligns with their professional conduct not to leave out available information and is essential to consider biodiversity data as part of the decisions*”, according to one participant from the United States. Participants also presented arguments opposing the notion that BIPD is not feasible in the already developed areas. As one participant from China stated, “*There are numerous examples that BIPD is implemented in the developed areas; for instance, in high-density cities, we can involve biodiversity vertically*.”

### 3.6 Post-sort questions

The top three suggestions put forth by participants regarding the involvement of BIPD in urban practice were concentrated across eight themes (Fig. 2 and 3): Data availability and methodology awareness, Public engagement and education promotion, Regulatory framework incorporation, Consensus building and synergy with other public interests, Best practice case studies and pilot projects, Multidisciplinary collaboration, Financing, and incentives and Others. The leading three are Data availability and methodology awareness, Public engagement and education promotion, and Regulatory framework incorporation, with the latter two being uniquely proposed across all four perspectives (Fig. 3). Therefore, we categorized these themes into four categories (Fig. 2): Knowledge-related, Engagement-related, Structure-related, and Others, with Knowledge-related constituting the top category, accounting for 41.7% of all responses (Fig.3). Notably, the suggestions of *Pluralists* and *Pragmatists* are relatively balanced across the first three categories (Fig. 2), while *Enthusiasts* exhibit a stronger preference for Knowledge-related suggestions (66.7%). Conversely, *Contextualists* provided no suggestions related to knowledge. This finding aligns with the Q-sort results, indicating that *Enthusiasts* are particularly passionate about incorporating biodiversity data, while *Contextualists* express a heightened sense of vulnerability regarding all propositions that are beyond their control and based on external or structural factors. Both *Pluralists* and *Pragmatists* suggested all three themes within the Knowledge-related category, with Data availability and methodology awareness being the most popular, while Multidisciplinary collaboration was the least favored.

**Fig.1:**
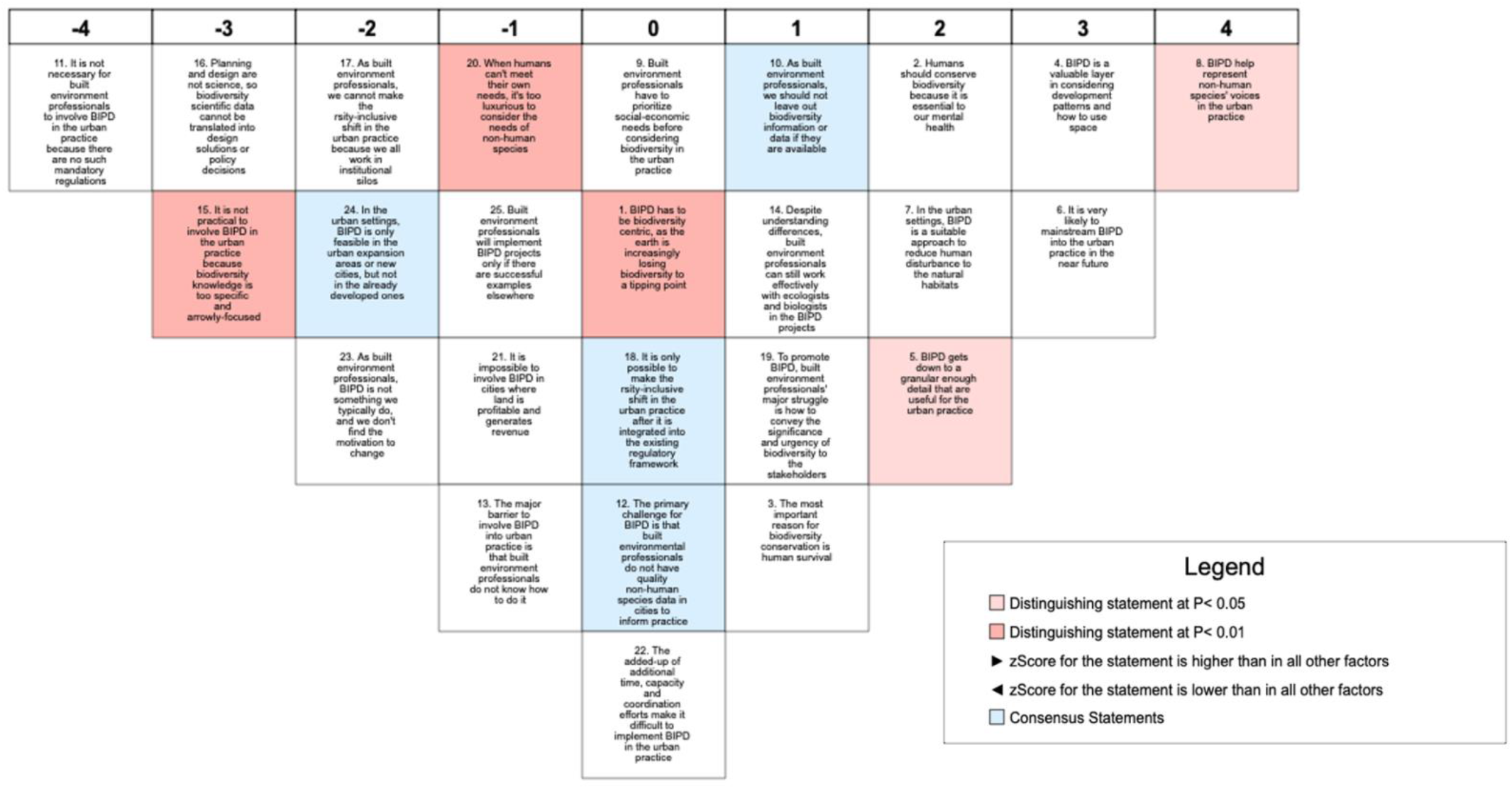
A Composite Ideal Sort aggregated by Ken-Q software by factor on a nine-column Q-Sort distribution board. Participants place one statement in each blank, and there is no difference in value based on the vertical order of statements in each column.

**Fig. 2:**
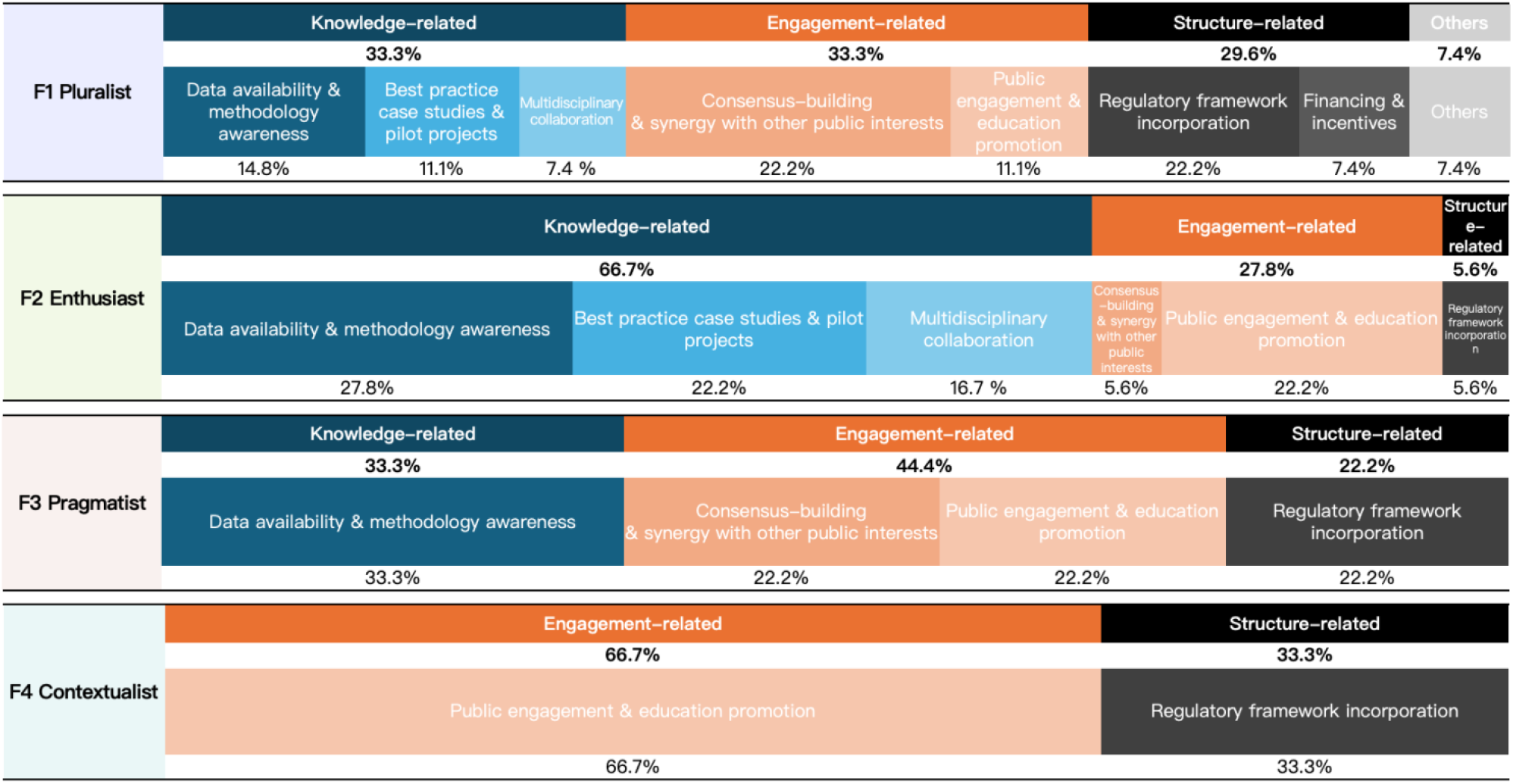
Bar chart of percentage by counts of the participants’ top three suggestions per factor. Suggestion categories are grouped in Knowledge-related (in blue), Engagement-related (in orange), Structure-related (in dark grey) and Others (in light grey).

**Fig. 3:**
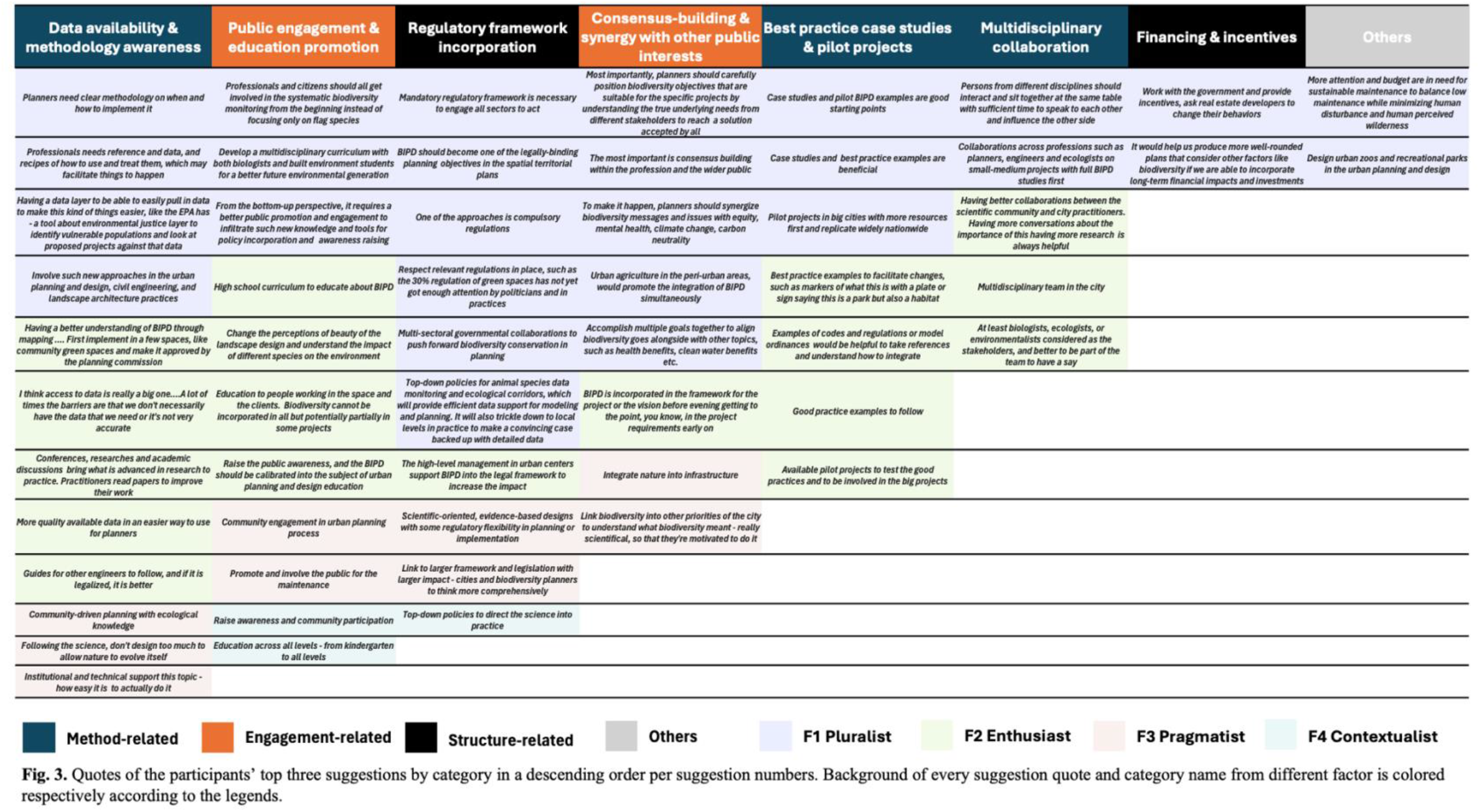
Quotes of the participants’ top three suggestions by category in descending order per suggestion numbers. Background of every suggestion quote and category name from different factor is colored respectively according to the legends.

There is a discernible trend toward increasing support for the statutory incorporation of BIPD, transitioning from “Not necessary” to “Statutory” (Table 4). This trend is evident across all groups, although the degree of support varies significantly. *Pluralists, Enthusiasts, and Contextualists* exhibit a gradual increase in support for integrating BIPD, demonstrating a clear preference for statutory incorporation. However, *Pragmatists* present a less clear pattern. Their responses indicate support for integrating BIPD while valuing diverse incorporation methods beyond legal frameworks. A participant from the US strongly agrees with the conceptual integration, stating that “*conceptual incorporation about biodiversity consideration is important if it is done well, … because they are rules of thumb that you can use that would make a huge difference compared to what we do now*.” Another Ethiopia participant also supports conceptual integration but feels neutral about integrating as plans or policies because “*indigenous residents have valuable conceptual insights into local wildlife and green spaces, (which) should be valued and incorporated*.”

**Table 4:** Participants’ intentions of incorporating BIPD into the legal framework by factors according to four five-point Likert scale selections

| Incorporation Level | Mean | Median | Standard Deviation | Minimum | Maximum |
| --- | --- | --- | --- | --- | --- |
| **F1 Pluralist (n=9)** |  |  |  |  |  |
| Not necessary | 1.2 | 1.0 | 0.4 | 1 | 2 |
| Conceptual | 1.6 | 1.0 | 0.8 | 1 | 3 |
| Non-statutory | 2.7 | 3.0 | 0.8 | 2 | 3 |
| Statutory | 4.4 | 4.0 | 0.5 | 4 | 5 |
| **F2 Enthusiast (n=6)** |  |  |  |  |  |
| Not necessary | 1.0 | 1.0 | 0.0 | 1 | 1 |
| Conceptual | 2.0 | 2.0 | 0.8 | 1 | 3 |
| Non-statutory | 2.0 | 2.0 | 0.6 | 1 | 3 |
| Statutory | 4.5 | 4.5 | 0.6 | 4 | 5 |
| **F3 Pragmatist (n=3)** |  |  |  |  |  |
| Not necessary | 1.0 | 1.0 | 0.0 | 1 | 1 |
| Conceptual | 3.7 | 4.0 | 2.1 | 1 | 5 |
| Non-statutory | 3.7 | 4.0 | 1.5 | 2 | 4 |
| Statutory | 4.7 | 5.0 | 0.6 | 4 | 5 |
| **F4 Contextualist (n=1)** |  |  |  |  |  |
| Not necessary | 1.0 | 1.0 | NaN | 1 | 1 |
| Conceptual | 3.0 | 3.0 | NaN | 3 | 3 |
| Non-statutory | 4.0 | 4.0 | NaN | 4 | 4 |
| Statutory | 5.0 | 5.0 | NaN | 5 | 5 |

Note: The four choices are defined as follows: Not necessary to incorporate, Conceptual incorporation is good enough, Non-statutory incorporation is sufficient, and Statutory incorporation is necessary. Likert scale: 1 = strongly disagree, 2 = disagree, 3 = neutral, 4 = agree, 5 = strongly agree. NaN indicates that the standard deviation cannot be calculated for a sample size of 1.

## Discussion

Our study provides well-informed and nuanced evidence to enhance understanding of the challenges and opportunities in realizing BIPD from the perspectives of BEPs. We engaged a diverse group of BEPs from geographically understudied regions in the Global South and professionals across sectors with socioeconomic expertise who have not typically been involved in this topic. Unlike previous studies that emphasize barriers arising from BEPs’ low-value priorities regarding ecology and their reluctant attitudes^39,42^, our findings are surprisingly positive considering the broad profile of BEPs examined. We identified three major challenges: the misalignment of value orientations between BEPs and their stakeholders, structurally imposed barriers, and a lack of basic awareness among BEPs about the existence of biodiversity data and methodologies. Our study identified potential opportunities that have not been fully explored in previous research regarding BEPs’ willingness and capacity to overcome certain challenges, while other challenges may require additional support, particularly from ecologists. We will first discuss these challenges and the perceived behavioral control of BEPs associated with them, along with BEPs’ coping mechanisms. This will be followed by a discussion of the potential opportunities forward, referencing the frameworks of the TPB^44^ and lock-in effects^41^.

Our study reveals that BEPs in both the Global North and South hold positive attitudes toward biodiversity despite their diverse value patterns associated with nature. These values resonate with the multiple ways in which individuals conceive a good quality of life, as discussed in the IPBES report^4,49^. For instance, *Enthusiasts* who support prioritizing biodiversity in all sorts of planning practices hold an ecocentric worldview, while the *Pluralist* recognizes the interconnectedness of systems, exhibiting a more pluricentric worldview. Regarding specific values, both *Enthusiasts* and *Contextualists* tend to acknowledge the intrinsic value of biodiversity, whereas *Pragmatists* appear to celebrate its instrumental value more. Despite these diverse value patterns, all BEPs favor BIPD and are willing to integrate it into their practices. This is evident from the consensus statement among BEPs that they will not overlook biodiversity information if it is made available and that they support the statutory incorporation of BIPD across perspectives.

However, BEPs perceive a disconnect between their ecological values and attitudes and the viewpoints of their stakeholders, over which they feel limited perceived behavioral control and seek assistance from additional data and methods beyond their current professional knowledge. All BEPs, except for *Pragmatists*, identify a major struggle in conveying the significance and urgency of biodiversity to stakeholders. In their view, the challenge is not due to BEPs’ reluctance or resistance, but rather a mismatch between BEPs’ ecological values and attitudes and those of their stakeholders – namely, residents, officials, clients, and others – acting as a significant barrier to the implementation of BIPD. To cope with this mismatch, BEPs discussed their attempts primarily in their engagement-related suggestions, including engaging the public, promoting education, and synergizing biodiversity with other public interests. Such coping approaches rested within their accustomed professional knowledge; however, BEPs also felt a sense of insufficiency when addressing such challenges by solely relying on their current knowledge. For instance, *Enthusiasts*, who experience a heightened struggle to convey the significance of biodiversity to their stakeholders, place greater hope in acquiring additional biodiversity-related knowledge, as evidenced by a high proportion (66.7%) of knowledge-related suggestions. These suggestions encompass a broad array of ways of accessing data and methods, case studies, and multidisciplinary collaborations. *Pluralists* made similar knowledge-related suggestions, but in a more balanced manner alongside other categories of suggestions, and they also emphasized the importance of sufficient details in the example methods of SDMs to help convey the significance of biodiversity while engaging with stakeholders.

Regarding the challenges associated with structural barriers, BEPs demonstrate varying degrees of perceived control. Unlike previous studies emphasizing inadequate legislation as a significant barrier^42^, BEPs in our study favor the statutory incorporation of BIPD while feeling a stronger sense of control over this barrier. This sense of control is supported by BEPs’ professional norms, as rules-in-use enable them to operate beyond the structural barriers of legislation, which exist as rules-in-form, to better serve the public interest^50^. Consequently, some BEPs, particularly *Pluralists* and *Enthusiasts*, generally view themselves as capable of transcending legislation and regulations, regarding this as an integral part of their role and professional ethical conduct in the decision-making process and in fostering a better urban future. This flexibility and potential for innovation reduce the degree of structural lock-in^44^ for them and reveal opportunities to act on BIPD involving BEPs’ expertise.

In contrast, BEPs’ perceived control over the structural-physical barriers^41^, such as land use conflicts and competing resources, is more limited with their current expertise alone. Both *Contextualists* and *Pluralists* mentioned their dilemma in balancing the competing agendas of biodiversity with the socioeconomic public interest, particularly in the Global South, when their stakeholders are often economically locked in^41^ and require innovative, tailored solutions. They either experience a sense of vulnerability beyond their control or address these competing agendas by prioritizing the socioeconomic needs of the public or postponing biodiversity considerations to the future. In this context, BEPs often feel they lack options. As a result, their actions may reflect a path dependency that prioritizes socioeconomic needs while downgrading biodiversity considerations, potentially attracting criticism for resistance to change with their socio-technical regime^42^. However, it seems likely that BEPs struggle more due to their inadequate capacity to engage stakeholders and collaboratively move towards BIPD than from their attitudinal resistance to change. For instance, upon learning about SDM examples for the first time, some *Pluralists* expressed that this data-rich evidence would help facilitate stakeholders’ understanding of the significance of biodiversity, demonstrating a strong interest in these data and methods and an expectation that such biodiversity evidence would assist in addressing the challenges of competing agendas, combined with their expertise in consensus building with stakeholders.

From the previous discussion, we can observe a pattern in which BEPs resort to additional biodiversity knowledge when they exhaust resources within their existing professional toolkits to address the challenges of misaligned values and structural-physical barriers. This is evidenced by the fact that they proposed the highest number of suggestions within the Knowledge-related category. Interestingly, BEPs did not endorse the statement that their lack of ecological knowledge is a major barrier in the Q-sort. One possible explanation for this contradiction may lie in BEPs’ lack of awareness of their own ecological knowledge deficits. Previous studies^16^ have pointed out BEPs’ lack of empirical ecological knowledge; however, a more pressing issue is that many BEPs may simply be unaware that such ecological data and methods exist. Without this foundational awareness of such knowledge, it may be challenging for them to accurately assess whether their lack of ecological knowledge constitutes a major barrier or not, let alone incorporate it into their practices. When presented with relevant data and methods, BEPs immediately express interest and perceive them as helpful, particularly in that the explicit and visual representations, supported by sufficient data, help speak to the sense and sensibility of the stakeholders so that they would care. Nevertheless, in addition to recognizing their usefulness, some *Pluralists*, particularly with previous experience of the SDM examples, voiced concern regarding the applicability of these tools, suggesting that currently available data and methods developed in urban ecology may not be readily actionable for prescribing real-world solutions. These findings reinforce existing research that emphasizes the knowledge gaps and disciplinary silos hindering BIPD^39, 40^ and further highlight the critical absence of basic awareness among many BEPs regarding the existence of ecological data and methodologies. Consequently, our findings emphasize the necessity of strategies that promote positive changes by empowering BEPs with a deeper understanding of advancements in ecological knowledge and improving the practical applicability of these methodological advancements from ecologists.

Having discussed the three identified challenges and their coping strategies, we find promising opportunities for realizing BIPD by learning from the perspectives of a broader profile of BEPs. First, BEPs exhibit positive attitudes toward biodiversity despite their varying value patterns associated with nature. Their professional norms facilitate adaptive changes by involving biodiversity in their practices. They are willing to overcome structural regulatory barriers and bridge gaps in values and attitudes with stakeholders through public engagement and consensus-building initiatives. If equipped with better biodiversity knowledge, particularly through evidence-based and readily applicable tools, BEPs may experience improved perceived control in communicating effectively with their stakeholders during these initiatives regarding competing structural-physical conflicts, facilitating a collaborative transition towards BIPD.

A consensus among all perspectives indicates that BEPs will not overlook biodiversity data if it is available to them. Specifically, BEPs are likely to act quickly when they are aware of advancements in biodiversity knowledge, informed on how to access such data, employ relevant methods, or understand whom to consult or collaborate with. *Pluralists* and *Enthusiasts* demonstrate a greater willingness to acquire more knowledge, proposing various Knowledge-related suggestions, including increasing data availability and methodological awareness, drawing inspiration from successful case studies, and pursuing collaborations with other disciplines.

Learning from BEPs’ perspectives, not all challenges are equally difficult. The more pressing barrier impeding BIPD lies in the fact that biodiversity data and methods have not yet been made available and applicable for BEPs. This echoes similar claims in related fields emphasizing disciplinary silos and knowledge gaps^16,26,39,42^. However, in addition to this, primary efforts should not only focus on raising awareness and understanding of biodiversity knowledge among BEPs, as emphasized by ecologists^16, 39^, but also on providing accessible biodiversity data and methods to BEPs, especially those urgently in need and those that are practically applicable. In our findings, for a broader profile of BEPs across sectors, professional backgrounds, and geographical regions globally, the vast majority have very limited awareness of the existence of biodiversity data, tools, and knowledge advances made by ecologists. Furthermore, those few BEPs with experience in BIPD expressed concern that the tools developed by ecologists are not easily applicable in urban planning and design practices. In this study, BEPs from various perspectives indicate that a key direction for the much-needed advances is the development of applicable evidence-based tools to facilitate effective communications in engaging with urban stakeholders for a collective transformative shift toward biodiverse cities.

This is hardly a single disciplinary effort. An imperative step forward is to collaboratively advance biodiversity knowledge and build multidisciplinary evidence^51,52^ to empower BEPs with accessible data and applicable tools. Our findings suggest that the common ground between the two disciplinary perspectives may already exist^40^. A noteworthy example is the *Pluralist* perspective identified in our study, which is widely shared among BEPs and aligns well with the value turn recently promoted within the conservation ecology community^4,52^, transitioning from “nature for itself” to “nature for people” ^53^. Another example is the *Enthusiast* perspective, characterized by a propensity for ecocentric values. These practitioners may radically approach a biodiverse urban future and exhibit greater openness to adapting new methods and collaborating with ecologists. Additionally, we have identified some novel research endeavors along this line in urban ecology, such as incorporating urban ecology data and methodological advances into the standard planning and design framework ^12,24^, balancing scientific measures and community meaningfulness in selecting target species and developing biodiversity objectives in an urban renewal project^37^. Still, further research is needed to better understand how to facilitate transdisciplinary knowledge advancement and awareness. For instance, further studies could explore the methods and tools that BEPs consider applicable in BIPD, strategies to transfer the knowledge advance and narrow gaps between urban ecologists and BEPs, and the role of biodiversity in setting objectives for BIPD.

In conclusion, our research challenges the notion that BEPs resist incorporating biodiversity into their practices by broadly exploring their viewpoints. Conversely, BEPs appreciate biodiversity and demonstrate both the desire and ability to overcome legislative structural barriers to implement BIPD. Insights from their perspectives reveal that the primary challenge hindering BIPD is the unavailability and inapplicability of biodiversity data and methodologies for BEPs. To fully leverage their potential and reshape urban environments, it is vital to equip BEPs with accessible biodiversity information and collaborate with them to create pragmatic, applicable solutions, especially in developing evidence-based tools for effective communication with urban stakeholders. This is particularly crucial in more challenging regions, such as the Global South, where contextualized strategies through stakeholder co-production are necessary. Transdisciplinary collaboration, with increased partnerships between ecologists and BEPs from diverse geographic areas and professional backgrounds, could elevate biodiversity considerations from advocacy to essential elements within comprehensive plans and urban renewal initiatives. This could lead to meaningful changes and potentially mainstream BIPD to transform cities into biodiversity-rich areas.

## Methods

### 2.1 Q methodology

We employ Q methodology, a mixed quantitative and qualitative research technique^54^, to illustrate BEPs’ typology of viewpoints on BIPD. This methodology has been increasingly utilized in conservation and sustainability research^55,56^. Q methodology employs factor analysis to statistically identify patterns of perspectives based on participants’ rank-sorting of items, allowing for interpretation and comparison of these perspective patterns^57^. While Q methodology is typically applied to understand viewpoints within a single geographical location, it can also be used to analyze viewpoints across multiple locations^54^.

To involve BEPs from both the Global North and South, we selected the United States, Spain, China, and Ethiopia as our case study countries, each characterized by distinctive levels of national urbanization rate^47^ and national GDP per capita, purchasing power parity (PPP)^48^. We also purposively sampled BEPs in these four countries to ensure diversity (Table 1), engaging those who may not typically have an interest in the topic of urban biodiversity across built-environment disciplines, including ecological-environmental and socioeconomic focus areas, and public, private, and non-profits sectors. We recruited 36 participants with professional experience ranging from 6 to over 30 years, yielding an average of 14.2 years of experience. Considering the diverse profile of the participants and the scarcity of BIPD practice, the vast majority of the recruited BEPs may not have knowledge or experience of BIPD.

To facilitate specific and concrete interviews, we briefly introduced BIPD and utilized examples of species distribution modeling and mapping^31,34^ to present a vivid illustration and enhance understanding. This research received ethical approval from the Universidad Politecnica de Madrid (UPM) (reference No. UAPGIAAESO-JAM-DATOS-20231114) before commencing, and all participants signed informed consent to participate.

### 2.2 Q statements

We initially conducted thirteen semi-structured interviews from December 2023 to February 2024 to develop a concourse of 340 statements^57^, reflecting BEPs’ intentions to incorporate BIPD in their practices. We inductively coded the interview transcripts using NVivo 12^58^ and categorized the statements into four themes: Environmental Concerns, Attitudes, Subjective Norms, and Perceived Behavioral Control (TPB)^44,46^. The last theme of TPB included Understanding Silos and Lock-in Effects^41^.

From this coded concourse, we selected 25 representative statements to match with the P-sample of 23 participants for Q-sorting interviews^59^. These statements were edited for clarity and relevance across countries and cultures while retaining the original meaning (Table 3). The statements were translated into Chinese and Spanish, and the translations were validated by both ecologists and BEPs native in these languages for use in China and Spain. For participants in Ethiopia, we communicated in English, having consulted their language preferences. English is widely used alongside their federal working language, Amharic, across all sectors in Ethiopia; for example, the nation’s Constitution is written in both English and Amharic^60^. We tested the Q-set with two respondents to ensure the statements were clear, comprehensive, and suitable to conduct for both face-to-face and online settings.

### 2.3 Conducting Q sorts

The Q sorts and post-sort interviews took place between March and April 2024. Each interview lasts between 60 and 90 minutes. We carried out all Q sorts using the Q-Tip platform (https://qtip.geography.wisc.edu/), a free online tool for conducting Q-method research developed by the University of Wisconsin-Madison. Participants were asked to sort the 25 statements onto a nine-column distribution board from Q-Tip (Fig. 1) based on their opinions regarding the extent to which they agreed or disagreed with each statement relative to others. During the post-sort interviews, we first discussed the participants’ sorts and then posted two follow-up questions. The first question aimed to capture BEPs’ proposed strategies by asking for their top three suggestions for approaches that they believe would help integrate BIPD into urban practice. The second question sought to gauge their intentions for change by having them select from a five-point Likert scale to rate how BIPD should be incorporated into the legal framework, ranging from “not necessary, ““conceptual,” “non-statutory” to “statutory” (Table 4).

### 2.4 Q analysis

Following standard analytical processes in Q Method^57^ and recommendations to minimize bias^61^, we analyzed the Q sorts using the R package ‘qmethod’ version 1.8.4.^62^ and Ken-Q Analysis software version 2.0.1^63^. After computing intercorrelations among individual Q sorts, we extracted four factors using Principal Components Analysis with eigenvalues greater than 1.77 and applied Humphrey’s rule to test the retention of these factors. The cumulative explanatory variance of these four factors was 70.8% (Table 2). We used the default Varimax rotation and retained the automatically flagged defining sorts for each factor, with the resulting minimum factor loading exceeding 0.60.

To aid in interpretation, we consulted the composite ideal sorts (Table 3 and Fig. 1), as aggregated by Ken-Q Analysis software^63^, which represent a shared viewpoint for each factor. Further, we employed NVivo 12^58^ to organize each factor’s post-sort interview transcripts and gain insights into participants’ perspectives through illustrative quotes.

## Acknowledgments

This research is funded by Jing Lu’s Marie Skłodowska-Curie Actions Grant No. 945139 (European Union’s Horizon 2020 Research and Innovative Programme) as part of her Ph.D. dissertation, as well as Fundación Tatiana Pérez de Guzmán el Bueno in Spain. The funder played no role in the study design, data collection, data analysis and interpretation, or this manuscript’s writing. We are grateful to Alicia Lopez, Jing Gan, Melissa Herliz, Tinsae Yimam, and Yonas Abate for their invaluable help in recruiting interview participants and all interviewees for sharing their valuable views and perspectives.

## Data Availability

All data generated or analyzed during this study are provided where possible in this paper, though individuxal responses and comments have not been provided to maintain confidentiality. The corresponding author can be reached for reasonable requests.

